# Granulocyte colony-stimulating factor protects against arthritogenic alphavirus pathogenesis in a type I IFN-dependent manner

**DOI:** 10.1101/2024.10.09.617470

**Authors:** Muddassar Hameed, Md Shakhawat Hossain, Andrea R. Daamen, Peter E. Lipsky, James Weger-Lucarelli

**Affiliations:** Department of Biomedical Sciences and Pathobiology, Virginia-Maryland College of Veterinary Medicine, Virginia Tech, Blacksburg, VA 24061, USA; Center for Zoonotic and Arthropod-borne Pathogens, Virginia Polytechnic Institute and State University, Blacksburg, VA 24061, USA; Department of Pathology & Immunology, Alvin J. Siteman Cancer Center, Washington University School of Medicine, St. Louis, MO 63110, USA; AMPEL BioSolutions LLC and the RILITE Research Institute, Charlottesville, VA, United States

## Abstract

Arthritogenic alphaviruses cause disease characterized by fever, rash, and incapacitating joint pain. Alphavirus infection stimulates robust inflammatory responses in infected hosts, leading to the upregulation of several cytokines, including granulocyte colony-stimulating factor (G-CSF). G-CSF is secreted by endothelial cells, fibroblasts, macrophages, and monocytes and binds to colony stimulating factor 3 receptor (CSF3R, also known as G-CSFR) on the surface of myeloid cells. G-CSFR signaling initiates proliferation, differentiation, and maturation of myeloid cells, especially neutrophils. Importantly, G-CSF has been found at high levels in both the acute and chronic phases of chikungunya disease; however, the role of G-CSF in arthritogenic alphavirus disease remains unexplored. Here, we sought to test the effect of G-CSF on chikungunya virus (CHIKV) and Mayaro virus (MAYV) infection using G-CSFR-deficient mice (G-CSFR^−/−^). We observed sustained weight loss in G-CSFR^−/−^ mice following viand MAYV infection compared to wild-type mice. Furthermore, G-CSFR^−/−^ mice had a significantly higher percentage of inflammatory monocytes and reduction in neutrophils throughout infection. The difference in weight loss in G-CSFR^−/−^ mice induced by alphavirus infection was corrected by blocking type I IFN signaling. In summary, these studies show that type I IFN signaling contributes to G-CSFR mediated control of arthritogenic alphavirus disease.

## Introduction

Arthritogenic alphaviruses, including chikungunya virus (CHIKV), Ross River virus (RRV), O’nyong nyong virus (ONNV), and Mayaro virus (MAYV) are significant public health threats (1, 2). These viruses are spread worldwide: CHIKV is endemic to Africa, Southeast Asia, and, more recently, the Caribbean and South America. RRV, ONNV, and MAYV are endemic to Australia, Africa, and South America, respectively (3). The disease caused by arthritogenic alphaviruses is characterized by high fever, rash, myositis, and arthritis, which can persist for years in roughly half of affected patients (4–9). However, there are no specific therapies available for the treatment of alphavirus arthritis except the use of anti-inflammatory drugs for symptomatic relief (10, 11). Thus, there is an urgent need to understand the immune mechanisms that control arthritogenic alphavirus disease outcomes in order to develop therapeutics.

During arthritogenic alphavirus infection, the virus replicates in fibroblasts, muscle satellite cells, macrophages, and other cells, initiating inflammatory responses (12, 13). This results in the influx of various immune cells, leading to tissue damage and the production of proinflammatory cytokines, including granulocyte colony-stimulating factor (G-CSF) (14–16). Previous studies have shown that G-CSF is upregulated in acute and chronic arthritogenic alphavirus-infected humans (17–20) and mice (21). G-CSF is a glycoprotein secreted from endothelial cells, fibroblasts, macrophages, monocytes, and bone marrow stromal cells (reviewed in (22, 23)). Its function is driven by binding to its cognate receptor, the G-CSF receptor (G-CSFR) (24–26), present on the surface of myeloid precursor cells and some non-immune cells. Binding initiates signal transduction and activation of cellular pathways that drive proliferation, differentiation, and maturation of granulocytes, especially neutrophils, (27, 28) which play an important role in arthritogenic alphavirus pathogenesis (29–31). Previous studies shows that higher influx of neutrophils into joint tissues during CHIKV infection is associated with worse disease outcomes in mice (30, 32). Furthermore, MyD88-dependent influx of neutrophils along with monocytes impairs lymph node B cell responses to CHIKV infection (29). Collectively, these data highlight the important role of G-CSF and neutrophils in arthritogenic alphavirus infection, which is still vastly understudied.

In this study, we investigated the contribution of G-CSF in the development of CHIKV and MAYV-induced disease. We infected G-CSFR knockout (G-CSFR^−/−^) (28) mice with CHIKV and MAYV and monitored disease development. We observed a sustained weight loss in infected G-CSFR^−/−^ animals compared to wild type (WT) control groups. Blood immune cell profiling showed that G-CSFR depletion led to a decrease in neutrophils and a significant increase in different monocyte populations throughout infection. Alphaviruses are highly sensitive to type I IFN restriction and monocytes are known to produce type I IFNs in response to CHIKV or RRV infection (33, 34). Therefore, we blocked IFN signaling, which corrected the weight loss differences observed between G-CSFR^−/−^ and WT mice. Future studies can be conducted to explore the role of G-CSF in mediating inflammatory responses during arthritogenic alphavirus infection, and the potential of targeting G-CSF or its receptor therapeutically.

## Materials and Methods

### Ethics statement

All experiments were conducted with the approval of Virginia Tech’s Institutional Animal Care & Use Committee (IACUC) under protocol number 24-060. Experiments using CHIKV and MAYV were performed in a BSL-3 or BSL-2 facility, respectively, in compliance with CDC and NIH guidelines and with approval from the Institutional Biosafety Committee (IBC) at Virginia Tech.

### Mice

C57BL/6J mice (strain# 000664) and colony stimulating factor 3 receptor gene knockout mice (B6.129X1 (Cg)-Csf3r^tm1Link/J^; herein referred to as G-CSFR^−/−^) (28) were purchased from The Jackson Laboratory at 6-8 weeks of age. Heterozygous G-CSFR^+/-^ mice were bred at Virginia Tech to obtain homozygous G-CSFR^−/−^ mice. Primers used to confirm homozygous colonies are presented in supplementary table S1. Homozygous G-CSFR^−/−^ mice were used to perform studies with CHIKV and MAYV infections. Mice were kept in groups of four or five animals per cage at ambient room temperature with *ad libitum* supply of food and water.

### Cell culture and viruses

Vero cells were obtained from the American Type Culture Collection (ATCC; Manassas, VA) and grown in Dulbecco’s Modified Eagle’s Medium (DMEM, Gibco) with 5% fetal bovine serum (FBS. Genesee), 1 mg/mL gentamicin (Thermo Fisher), 1% non-essential amino acids (NEAA, Sigma) and 25 mM HEPES buffer (Genesee) at 37°C with 5% CO_2_. CHIKV SL-15649 (ECSA lineage), rescued from an infectious clone, was a gift from Dr. Mark Heise (the University of North Carolina at Chapel Hill (35), and MAYV strain TRVL 4675 was derived from an infectious clone (36, 37). Virus titers were determined by plaque assay as previously described (38).

### Mice infection

Mice were injected in both hind footpads with 10^2^ PFU or 10^5^ PFU of CHIKV SL-15649 or 10^4^ PFU of MAYV in 50 μL viral diluent in each foot (39, 40). All virus dilutions were made in RPMI-1640 media with 10 mM HEPES and 2% FBS, hereafter referred to as viral diluent. Mice were monitored for disease development following infection through daily weighing, and footpad swelling was measured using a digital caliper. Blood was collected via submandibular bleed for serum isolation to determine viremia and cytokine levels. At sixteen or twenty-one-days post-infection (dpi), mice were euthanized, and blood were collected to isolate immune cells.

### Luminex assay and ELISA

We quantified G-CSF levels in the serum of mock– and CHIKV/MAYV-infected animals using the mouse Luminex XL cytokine assay (bio-techne) and the G-CSF DuoSet ELISA kit (Catalog# DY414-05, R&D Systems) according to the manufacturer’s instructions. A standard curve was generated using the optical density values of the standards, which were used to estimate the cytokine levels in each sample.

### Type I IFN signaling blockage

For *in vivo* interferon-α/β receptor (IFNAR) blockade, C57BL/6J and G-CSFR^−/−^ mice were intraperitoneally (i.p.) inoculated with 0.1 mg of MAR1–5A3, anti-IFNAR antibody (Leinco Technologies, Fenton, MO) (41) one day prior to infection. Mice were then inoculated with 10^2^ PFU of CHIKV SL-15649 in 50 μL of viral diluent in both hind feet and monitored for disease development until 21 dpi.

### Mouse blood immune cell isolation

Mouse blood leukocytes were isolated using Mono-Poly resolving medium (M-P M, MP Bio, Cat. No. 091698049) according to the manufacturer’s instructions. Briefly, blood was mixed with an equal volume of PBS and layered slowly onto M-P M followed by centrifugation at 300 × *g* for 30 min in a swinging bucket rotor at room temperature (20–25°C). We collected cell layers between the plasma and M-P M to isolate leukocytes and added them to a 15 mL conical tube containing 10 mL of cold 10% FBS containing RPMI-1640 (RPMI-10). Cells were spun at 500 × *g* for 5 min at 4°C and used for flow cytometry.

### Flow cytometry

Single cell suspensions were washed with PBS and resuspended in 100 μL Zombie aqua cell viability dye solution (1:400 prepared in PBS, Cat. No. 423101, BioLegend) and incubated at room temperature for 15-30 minutes. 200 μL flow cytometry staining (FACS) buffer (PBS containing 2% FBS) was added and centrifuged at 500 × *g* for 5 min at 4°C. The resulting cell pellet was resuspended in FACS buffer with 0.5 mg/mL rat anti-mouse CD16/CD32 Fc block (Cat. No. 553142, BD Biosciences) and incubated for 15 min on ice to block Fc receptors. For extracellular staining, a combined antibody solution was prepared in FACS buffer with fluorophore-conjugated antibodies: anti-mouse Alexa fluor 700 CD45 (Cat. No. 103128, BioLegend), anti-mouse PerCP/Cyanine 5.5 CD11b (Cat. No. 101227, BioLegend), anti-mouse APC Ly6G (Cat. No. 127614, BioLegend), anti-mouse PE Ly6C (Cat. No. 128007, BioLegend), anti-mouse PE CD3 (Cat. No. 100206, BioLegend), anti-mouse PerCP/Cyanine 5.5 CD4 (Cat. No. 116012, BioLegend), and anti-mouse FITC CD8a (Cat. No. 100706, BioLegend), and APC CD19 (Cat. No. 152410, BioLegend). 100 μL antibody cocktail was added to the single cell suspension, mixed, and incubated for 30 min on ice. Cells were washed with FACS buffer twice, and 100 μL 4% formalin (Thermo Fisher Scientific, Ref. No. 28908) was added to fix the cells. After 15 min incubation at room temperature, cells were washed with FACS buffer, resuspended in 100-200 μL PBS, and covered with aluminum foil before flow cytometry analysis. For each antibody, single color controls were run with Ultracomp ebeads (Cat. No. 01-2222-42, Thermo Fisher Scientific). The stained cells were analyzed using the FACSAria Fusion Flow cytometer (BD Biosciences).

### Statistical analysis

Statistical comparisons were performed using GraphPad Prism version 9. Data are presented as mean ± standard deviation. The statistical tests used to analyze data are described in figure legends.

## Results

### 1.1 G-CSF is upregulated in response to CHIKV and MAYV infection in C57BL/6J mice

CHIKV and MAYV produce disease in C57BL/6J mice similar to outcomes in humans (39). G-CSF is elevated in humans infected with arthritogenic alphavirus (17–20) and mice (21). To assess G-CSF levels following arthritogenic alphavirus infection, we infected mice with CHIKV and MAYV and collected blood at 2 and 7 dpi, the peak of viremia and footpad swelling, respectively (39). At 2 dpi, G-CSF levels increased 2.5-fold following CHIKV (*p=0.0079*) or 2-fold (*p=0.0001*) following MAYV infection (Fig. 1A-B) compared to mock-inoculated controls. To assess whether G-CSF remained elevated later in infection, we measured G-CSF levels in the blood at 7 dpi. Similarly, G-CSF levels were significantly higher at 7 dpi in CHIKV (*p=0.01*) and MAYV (*p=0.04*) infected mice, respectively, compared to mock-infected groups (Fig. 1C-D). This increase in G-CSF levels following CHIKV and MAYV infection in mice is consistent with reports in humans infected with arthritogenic alphaviruses (17–20).

**Figure 1.**
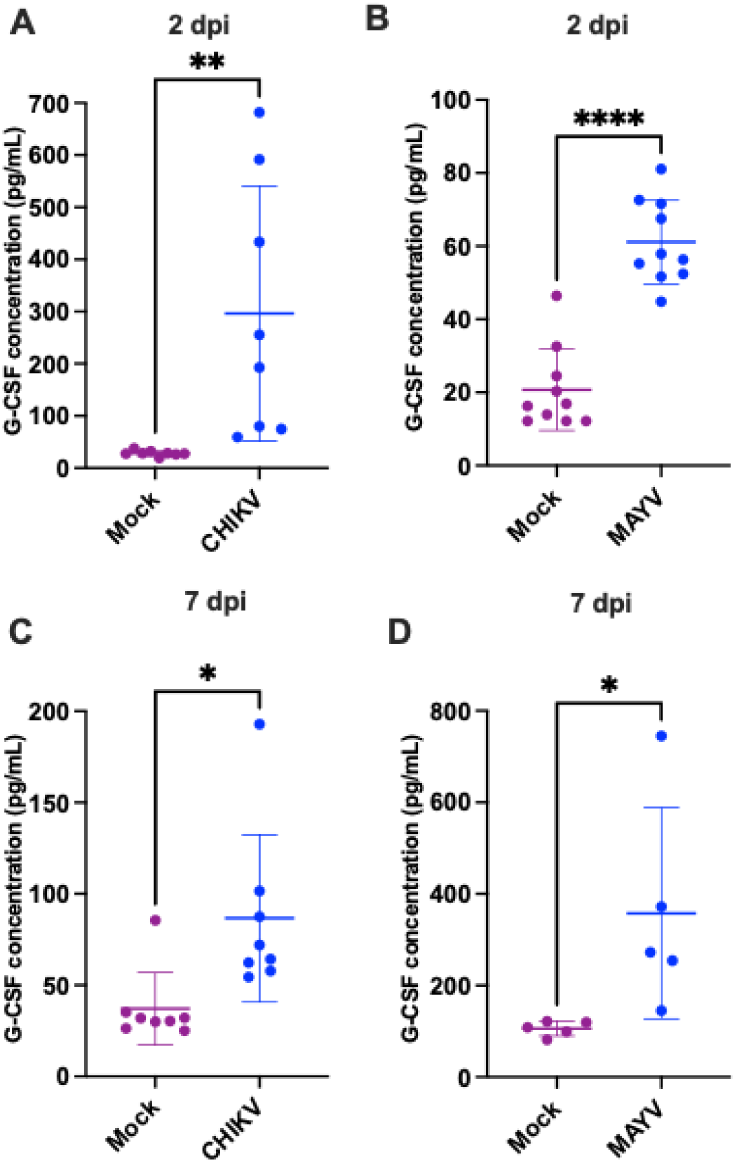
G-CSF is upregulated in response to CHIKV and MAYV infection in C57BL/6J mice. 6–8-week-old C57BL/6J mice were inoculated with viral diluent (mock), 10^5^ PFU of CHIKV strain SL-15649, or 10^4^ PFU of MAYV strain TRVL 4675 in each hind footpad, and blood was collected at 2 and 7 dpi for serum isolation. G-CSF levels were measured by ELISA and Luminex assay at 2 dpi (**A, B**) and by ELISA at 7 dpi (**C, D**), presented in picograms per milliliter of serum. The error bars represent the standard deviation, horizontal bars indicate mean values, and asterisks indicate statistical differences. Statistical analysis was done using an unpaired t-test; *, p < 0.05; **, p < 0.01; ****, p < 0.0001.

### 1.2 Lack of G-CSF signaling causes sustained weight loss in mice infected with arthritogenic alphaviruses

G-CSFR is present on the surface of granulocyte precursors, initiating signal transduction and activation of cellular pathways that drive proliferation, differentiation, and maturation of granulocytes, especially neutrophils (27, 28). To assess the impact of G-CSF signaling on arthritogenic alphavirus disease outcomes, we inoculated mice lacking the G-CSFR (G-CSFR^−/−^) (28). Following CHIKV infection, we observed a sustained weight loss in G-CSFR^−/−^ mice compared to WT mice until 21 dpi, when the study was terminated (Fig. 2A). CHIKV-infected G-CSFR^−/−^ mice had a trend toward higher footpad swelling at 6 dpi but there was no statistical difference (Fig. 2B). No difference was observed in virus replication at 2 dpi (Fig. 2C, Fig. S1). MAYV-infected G-CSFR^−/−^ mice showed a similar sustained weight loss compared to the WT group until 16 dpi (Fig. 2D). No difference was observed in footpad swelling and virus replication (Fig. 2E, 2F, Fig. S1). These data suggest that G-CSFR deficiency restricts the ability to recover from systemic arthritogenic alphavirus disease.

**Figure 2.**
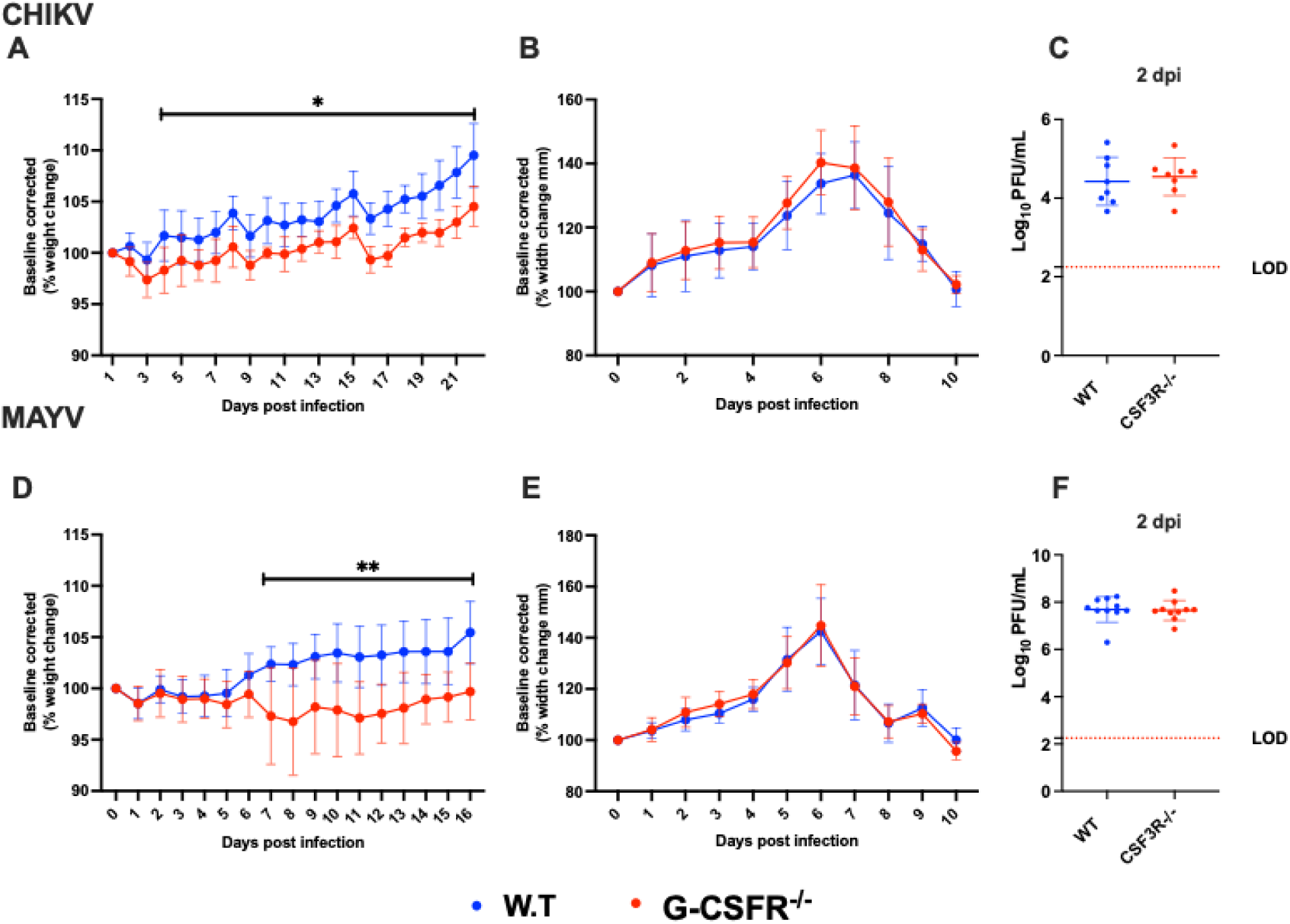
G-CSF receptor depletion cause sustained weight loss in mice infected with arthritogenic alphaviruses. C57BL/6J and G-CSFR^−/−^ mice were inoculated with viral diluent (mock), 10^5^ PFU of CHIKV strain SL-15649 (n=8), or 10^4^ PFU of MAYV strain TRVL 4675 (n=10) in each hind footpad and monitored for disease development until 21 and 16 dpi, respectively. **A-C.** Weight loss (**A**), footpad swelling (**B**), and the development of viremia (**C**) were determined after CHIKV infection. **D-F.** Weight loss (**D**), footpad swelling (**E**), and virus replication (**F**) were measured following MAYV infection. Weight loss and footpad swelling were analyzed using multiple unpaired t-tests with the Holm-Sidak method for multiple comparisons, and viremia data was analyzed by unpaired t-test with Welch’s correction. The error bars represent the standard deviation, bars indicate mean values, dotted line represents the limit of detection, and asterisks indicate statistical differences. The level of significance represented is as follows *p<0.05, **p<0.01.

### 1.3 Lack of G-CSF signaling leads to the systemic reduction of neutrophils and increase in monocytes during arthritogenic alphavirus infection

Next, we monitored the impact of the G-CSFR deficiency on the immune cell profile throughout the course of infection. We isolated blood leukocytes at various points following CHIKV– and MAYV-infection and profiled immune cell populations using flow cytometry. As expected, G-CSFR deficiency led to a significant reduction in the proportion of neutrophils in blood compared to WT mice during CHIKV and MAYV infection (Fig. 3A and 3D). We also observed a significantly higher percentage of anti-inflammatory monocytes in the blood of CHIKV-infected G-CSFR^−/−^ animals until 21 dpi (Fig. 3B). There was no difference in pro-inflammatory monocytes between G-CSFR^−/−^ and WT mice 2 dpi with CHIKV; however, differences emerged at 7 and 14 dpi, with pro-inflammatory monocytes increasing in proportion in G-CSFR^−/−^ mice before returning to baseline at 21 dpi (Fig. 3C). Similarly, the percentage of anti-, and pro-inflammatory monocytes were higher in G-CSFR^−/−^ mice throughout MAYV infection (Fig. 3E and 3F). These data demonstrate that G-CSFR^−/−^ deficiency leads to substantial differences in myeloid cell populations in the blood following arthritogenic alphavirus infection.

**Figure 3.**
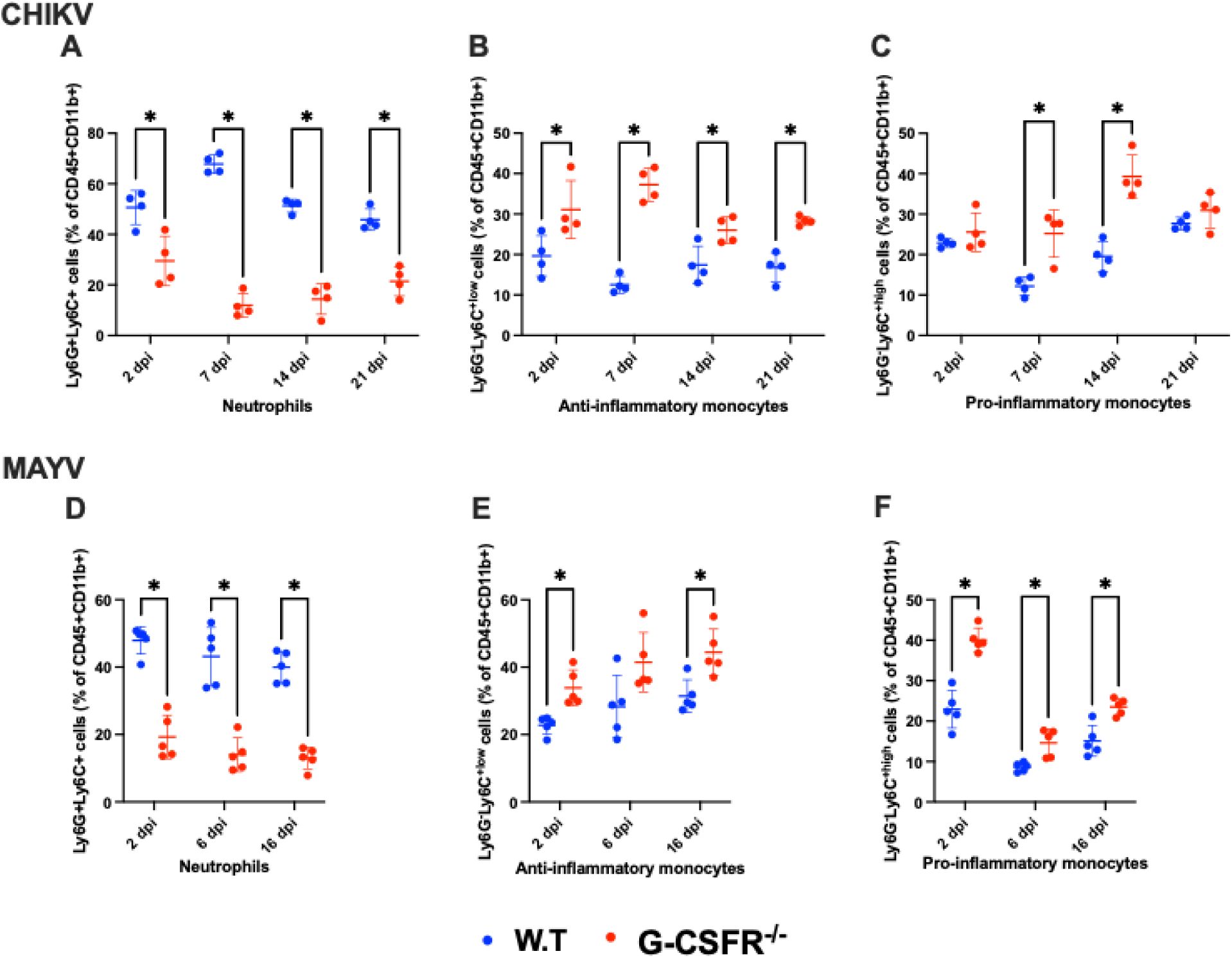
Lack of G-CSF function leads to the systemic reduction of neutrophils and increase in inflammatory monocytes during arthritogenic alphavirus infection. C57BL/6J and G-CSFR^−/−^ mice were inoculated with viral diluent (mock), 10^5^ PFU of CHIKV strain SL-15649 (n=4), or 10^4^ PFU of MAYV strain TRVL 4675 (n=5) in each hind footpad and blood was collected at the indicated timepoints. Leukocytes were isolated from the blood and subjected to flow cytometry. **A-F.** Dot plots presenting the percentage of neutrophils (CD45^+^CD11b^+^Ly6G^+^Ly6C^+^) (**A/D**), anti-inflammatory monocytes (CD45^+^CD11b^+^Ly6G^-^ Ly6C^low^) (**B/E**), and pro-inflammatory monocytes (CD45^+^CD11b^+^Ly6G^-^Ly6C^high^) (**C/F**) in WT and G-CSFR^−/−^ groups during CHIKV and MAYV infection. Data were analyzed using multiple unpaired t-tests and corrected with the Holm-Sidak method for multiple comparisons. The error bars represent the standard deviation, bars indicate mean values, and asterisks indicate statistical differences. The level of significance represented as follows *p < 0.05.

We also determined the effect of G-CSFR^−/−^ deficiency on B and T cell dynamics during arthritogenic alphavirus infection. We observed no impact on CD4 T cells in CHIKV– and MAYV-infected animals (Fig. S2A/D). CD8 T cells were similar between the groups except at 7 dpi following CHIKV infection, where the percentage of these cells was higher in G-CSFR^−/−^ mice; a similar difference was observed following MAYV infection (Fig. S2B/E). In contrast, B cells were higher in G-CSFR^−/−^ mice at 6/7 dpi following CHIKV and MAYV infection (Fig. S2C and S2F). These data demonstrate that G-CSFR^−/−^ deficiency has minimal impacts on the proportion of T cells during infection but leads to a significant increase in B cells at 6-7 dpi.

### 1.4 The impact of G-CSF on disease recovery depends on type I interferon signaling during arthritogenic alphavirus infection

Alphaviruses are highly sensitive to type I IFN restriction, and mice deficient in IFNAR-signaling are highly susceptible to disease (30, 42). Monocytes are highly enriched in G-CSFR^−/−^ mice (Fig. 3) following CHIKV and MAYV infection and are known to produce type I IFNs in response to CHIKV or RRV infection (33, 34). Thus, we hypothesized that type I IFN might be responsible for worse systemic disease in G-CSFR^−/−^ mice, and blockade of type I IFN signaling might reverse disease outcomes in G-CSFR^−/−^ mice. To test this, we injected MAR1-5A3 antibody (41) to block IFNAR signaling one day prior to CHIKV infection. We observed a similar pattern of weight loss in CHIKV-infected WT and G-CSFR^−/−^ groups, indicating that worse systemic disease observed in G-CSFR^−/−^ mice is mediated by type I IFN signaling (Fig. 4A). We observed no differences in footpad swelling (Fig. 4B) or virus replication between groups (Fig. 4C). Overall, these data highlight that G-CSF plays an important role in control of systemic alphavirus disease, which is dependent on type I IFN signaling.

**Figure 4.**
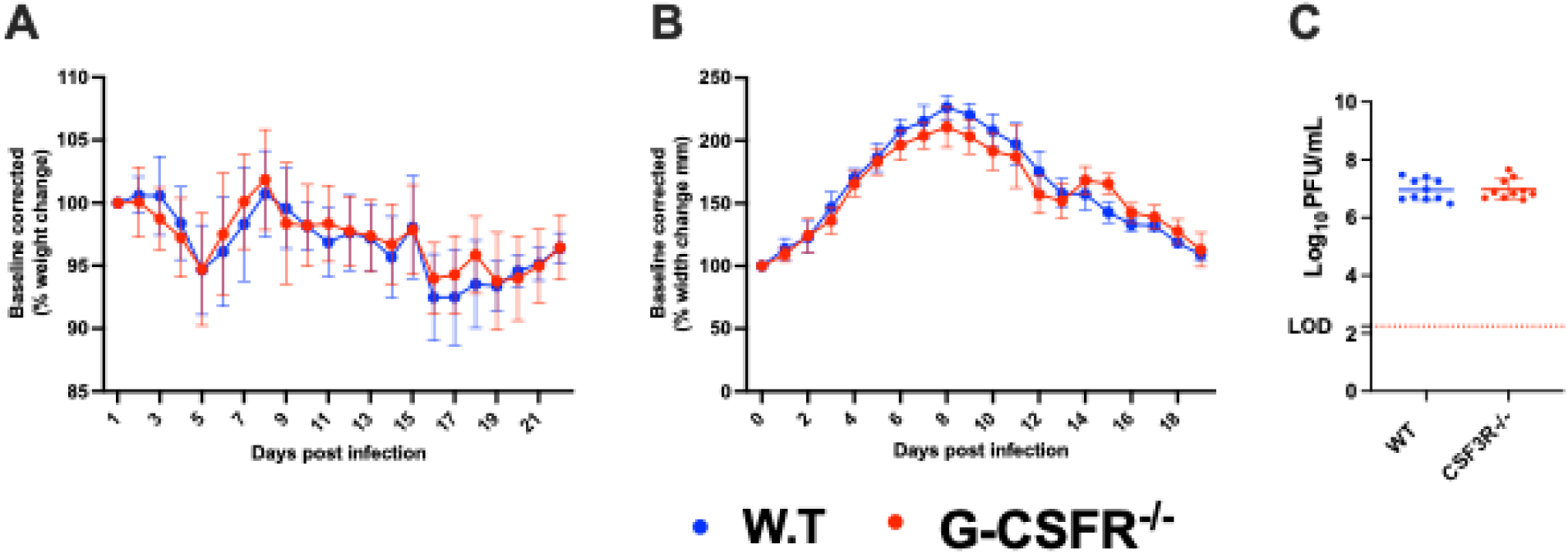
The impact of G-CSF on disease recovery depends on type I interferons during arthritogenic alphaviruses. C57BL/6J and G-CSFR^−/−^ mice were injected with 0.1 mg MAR1-5A3 antibody 1 day before infection to block type I IFN signaling. Mice were inoculated with 10^2^ PFU CHIKV strain SL-15649 in each hind footpad and monitored for disease development until 21 dpi (n=10). **A-C.** Weight loss (**A**), footpad swelling (**B**), and virus replication (**C**) was measured following CHIKV infection. Weight loss and footpad swelling were analyzed using multiple unpaired t-tests with the Holm-Sidak method for multiple comparisons, and viremia data was analyzed by unpaired t-test with Welch’s correction. The error bars represent the standard deviation, bars indicate mean values, dotted line represents the limit of detection, and asterisks indicate statistical differences.

## Discussion

Arthritogenic alphavirus infection causes acute and chronic disease characterized by fever, skin rash, myalgia, and debilitating joint pain. In the infected vertebrate host, inflammatory responses are activated in target tissues such as muscles and joints, leading to the up-regulation of several proinflammatory cytokines, including G-CSF (17–20). Here, we explored G-CSF’s role in arthritogenic alphavirus disease. We infected G-CSF receptor deficient mice and observed that CHIKV and MAYV infection cause sustained weight loss compared to WT mice. Blocking type I IFN signaling in G-CSFR^−/−^ mice through anti-IFNAR1 antibody injection reversed this phenotype. Furthermore, we observed a higher percentage of monocytes in CHIKV– and MAYV-infected G-CSFR^−/−^ mice compared to WT animals. Together, these data suggest that G-CSFR-signaling controls systemic arthritogenic alphavirus disease through type I IFN signaling.

G-CSF is a glycoprotein secreted by endothelial cells, fibroblasts, macrophages, monocytes and bone marrow stromal cells and upregulated in acute and chronic arthritogenic alphavirus-infected humans (17–20) and mice (21). Therefore, we hypothesized that G-CSF contributes to arthritogenic alphavirus disease. To test this, we inoculated G-CSFR^−/−^ mice (28) with CHIKV and MAYV and monitored disease outcomes, viral replication, and immune cell dynamics. We observed a sustained delay in weight gain following CHIKV and MAYV infection in G-CSFR^−/−^ mice compared to WT mice. No differences were observed in footpad swelling or systemic virus replication between G-CSFR^−/−^ and WT mice. As expected, we observed that G-CSFR^−/−^ mice had a significant reduction in the proportion of neutrophils in the blood compared to WT mice during CHIKV and MAYV infection (27, 28). Neutrophil infiltration has been implicated in more severe footpad swelling following CHIKV infection (32); however, depletion of neutrophils in WT mice has no impact on disease outcome (29). Neutrophils were replaced in G-CSFR^−/−^ mice with both anti-(Ly6C^low^) and pro-inflammatory (Ly6C^high^) monocytes. Recently, it has been reported that Ly6C^+^ monocytes facilitate alphavirus infection at the initial infection site, which promotes more rapid spread into circulation (43). Furthermore, Ly6C^high^ monocyte recruitment to the draining lymph nodes during CHIKV infection impairs the virus-specific B cell responses by virtue of their ability to produce nitric oxide (29). The increase in Ly6C^+^ monocytes in the absence of G-CSFR may have led to a dysregulated adaptive immune response, possibly contributing to sustained weight loss observed in G-CSFR^−/−^ mice. In addition, G-CSF-stimulated neutrophils may serve to control monocyte activation in blood during arthritogenic alphavirus infection as neutrophils-induced immune modulation has been reported previously (44–47). These data suggest that G-CSF serves to maintain neutrophil populations during alphavirus infection, potentially keeping other immune cells in check. Further investigation is required to explore the complex effect of G-CSF on neutrophil and other immune cell’s functionality during alphavirus infection.

Next, we aimed to explore the mechanism by which G-CSF contributes to controlling systemic arthritogenic alphavirus disease. In previous studies, it has been reported that dysregulated type I IFNs can lead to poor disease recovery from virus infection (48–51). Therefore, we hypothesized that G-CSF signaling might contribute to the regulation of type I IFN responses in alphavirus infected animals, mediating the control of systemic disease. When we blocked type I IFN signaling by blocking IFNAR1, we observed a similar pattern of weight loss in CHIKV-infected WT and G-CSFR^−/−^ mice. These findings highlight that G-CSF may provide a protective role in WT mice through the regulation of type I IFNs.

Future experiments are needed to explore G-CSF’s impact on the functionality of specific immune cell populations, including but not limited to neutrophils and monocytes. Furthermore, the role neutrophil’s play in systemic disease should be assessed since most studies have focused on footpad swelling. Finally, G-CSF’s potential as a therapeutic to reduce systemic arthritogenic alphavirus disease should be tested in future studies.

In summary, our study sheds light on the role of G-CSF signaling in CHIKV and MAYV pathogenesis. Our findings show that G-CSF signaling is crucial for controlling systemic disease during alphavirus infection. G-CSFR deficiency led to a marked reduction in neutrophils during infection, which were replaced by both anti– and pro-inflammatory monocytes and B cells in blood. Additionally, the effect of G-CSFR deficiency on weight loss was found to be dependent on type I interferon signaling. Given these insights, further exploration of G-CSF’s role at the molecular level in arthritogenic alphavirus disease is warranted. This could pave the way for its potential development as a therapeutic target for both acute and chronic forms of the disease.

## Acknowledgements

We are grateful to Melissa Makris for assisting with flow cytometry analysis. This work was supported by NIAID R21AI153919-01 awarded to J.W-L.

