## Supplementary figure 1 for "Granulocyte colony-stimulating factor protects against arthritogenic alphavirus pathogenesis in a type I IFN-dependent manner"

C57BL/6J and G-CSFR^-/-^ mice were infected with viral diluent (mock) or 10^5^ PFU of CHIKV strain SL-15649 and 10^4^ PFU of MAYV strain TRVL 4675 in each hind footpad and serum was collected at 1, 2, 3, and 4 dpi to determine the development of viremia. Data were analyzed using multiple unpaired t-tests. The error bars represent the standard deviation, bars indicate mean values, and dotted line represents the limit of detection.

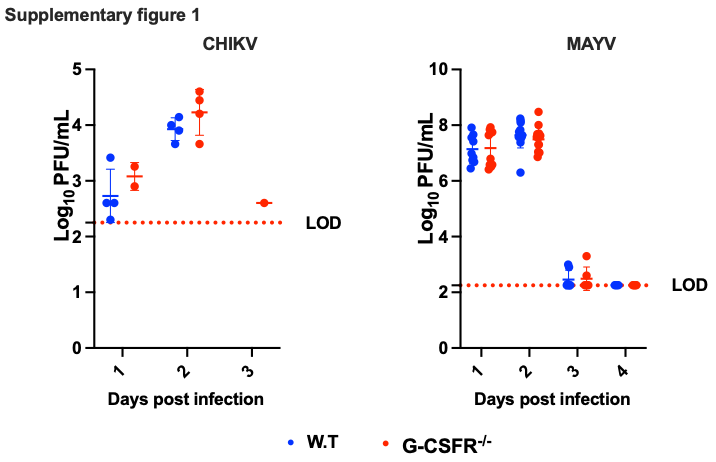
