## Supplementary figure 2 for "Granulocyte colony-stimulating factor protects against arthritogenic alphavirus pathogenesis in a type I IFN-dependent manner"

**Blood immune cell profiling during peak viremia and severe disease symptoms shows no considerable effect of lack of G-CSF function on adaptive immune cells during arthritogenic alphavirus infection.**

C57BL/6J and G-CSFR^-/-^ mice were infected with viral diluent (mock) or 10^5^ PFU of CHIKV strain SL-15649 and 10^4^ PFU of MAYV strain TRVL 4675 in each hind footpad and bled at 2, 6/7, 14, and 16/21 dpi. Leukocytes were isolated using mono-poly medium and subjected to flow cytometry. **A-F** Dot plots present the percentage changes in CD4 T cells (CD45^+^CD3^+^CD4^+^) (**A &D**), CD8 T cells (CD45^+^CD3^+^CD8^+^) (**B&E**), and B cells (CD45^+^CD3^-^CD19^+^). Immune cell percentage data were analyzed using multiple unpaired t-tests and corrected with the Holm-Sidak method for multiple comparisons. The error bars represent the standard deviation, bars indicate mean values, and asterisks indicate statistical differences. The level of significance represented as follows *p < 0.05, (n=4 or 5 each group).

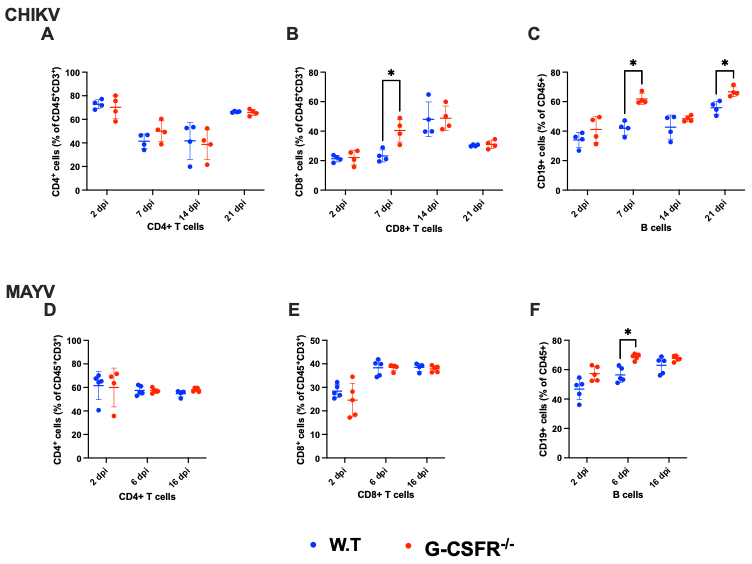
