## Supplementary table 1 for "Granulocyte colony-stimulating factor protects against arthritogenic alphavirus pathogenesis in a type I IFN-dependent manner"

List of RT-qPCR primers used for this study to perform the genotyping for G-CSFR^-/-^ mouse.

| Primer |  | Sequence 5' → 3' |
| --- | --- | --- |
|  | Common | ACATAAGCCTGTGGGAAGG |
|  | Wild type Reverse | GCTGGTTCTCCACTCATTTG |
|  | Mutant Reverse | CTCCAGACTGCCTTGGGAAAA |
